# mRNA Therapy for Alport Syndrome

**DOI:** 10.64898/2026.01.20.700554

**Authors:** Brian J. Parrett, Michael A. Barry

## Abstract

Alport syndrome is caused by mutations in the type IV collagen genes *COL4A3, COL4A4,* and *COL4A5* which are expressed in podocytes in the glomerulus of the kidney. Mutation of these genes disrupts blood filtration by the kidney leading to proteinuria and kidney failure. There is no cure for Alport Syndrome. Current therapies do not treat the genetic cause of the disease, but instead aim to delay the disease by reducing blood pressure in the kidney. Here we tested the ability of mRNA therapy to treat the disease. Mice with X-linked Alport syndrome (XLAS) due to mutation of *Col4A5* were treated intravenously with lipid nanoparticles (LNPs) carrying three mRNAs encoding *COL4A3, COL4A4,* and *COL4A5* to produce trimeric collagen IV repair proteins. This intravenous mRNA therapy significantly reduced proteinuria and blood urea nitrogen. Protection against the syndrome was maintained provided that LNP-mRNA injections were continued. However, efficacy was lost when therapy was terminated. These data provide proof of principle to apply a genetic mRNA therapy to halt progression of Alport syndrome in a mouse model of XLAS. These data also demonstrate that damage to the kidney filtration barrier can be leveraged to deliver large molecular therapies to podocytes and other cells within the kidney to treat Alport syndrome and other kidney genetic diseases.

## Introduction

Alport syndrome (AS) is a chronic genetic kidney disease with a wide range of disease severity. Clinical manifestations include hematuria, proteinuria, hearing loss, eye abnormalities, and kidney failure in severe cases (**Figure 1** and ^1^). Alport syndrome is caused by mutations in the type IV collagen alpha chain genes *COL4A3, COL4A4,* or *COL4A5.*^2^ The incidence of Alport syndrome ranges from 1:5000 to 1:53,000 live births.^3,4^ However more recent studies have indicated that the prevalence of potentially pathogenic variants can be as high as 1:106 or 1:2320 for *COL4A3/COL4A4* and *COL4A5,* respectively in the general human population.^5^ Other forms of kidney disease have also been found to be associated with mutations in these *COL4* genes, causing experts in the field to suggest reclassifying related nephropathies to AS to more appropriately address and manage the disease.^6,7^

**Figure 1:**
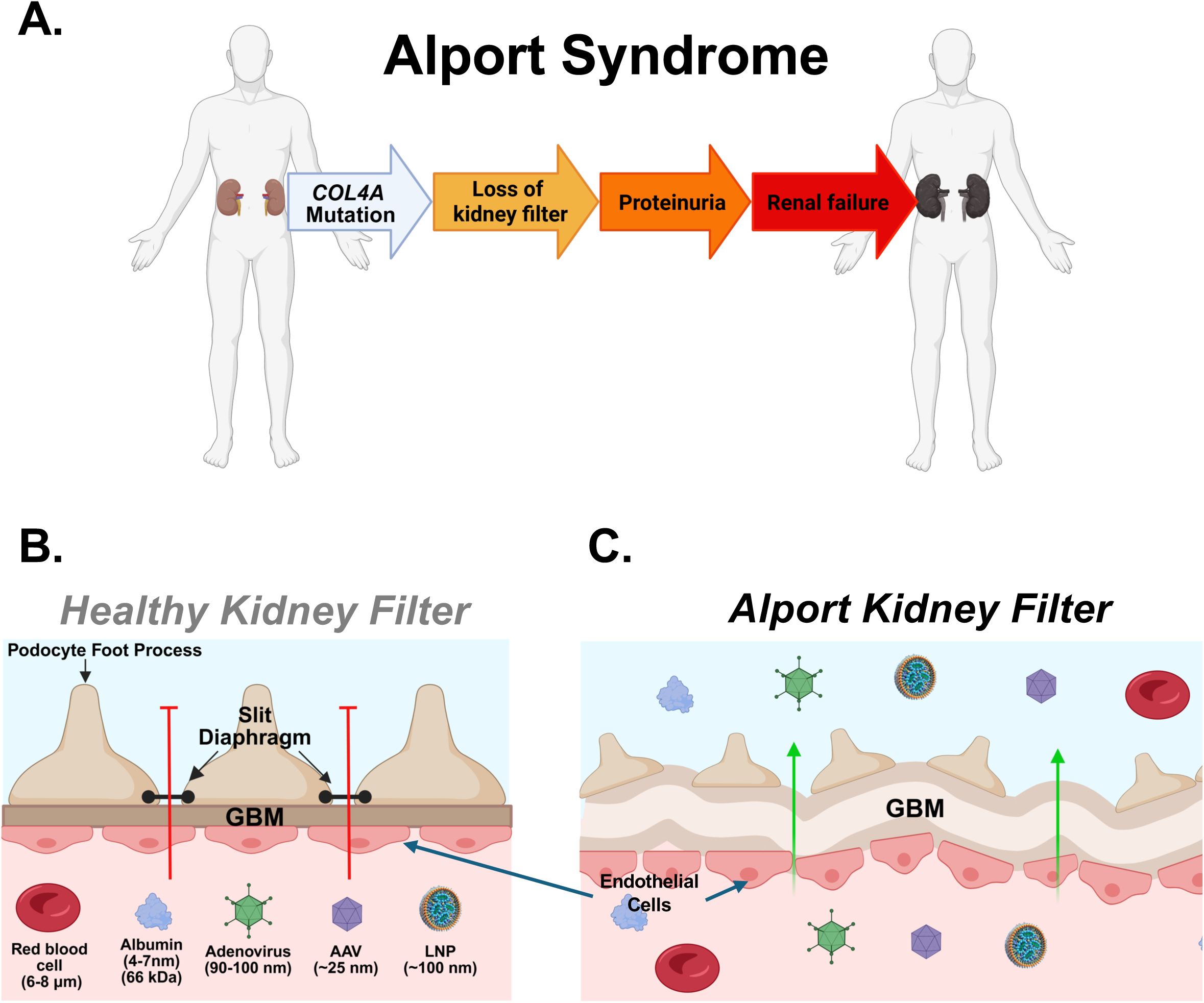
Visual representation of glomerular filtration barrier. Visual representation of the hypothesis that GFB permeability may increase molecular therapy vector access to the glomerulus. Healthy glomerular basement membranes (GBM) restricting passage of large particles (left panel). GBM damage in Alport syndrome allows larger particles to pass through the GBM, possibly increasing vector transfection in the glomerulus (right panel). AAV, Adeno-associated virus; LNP, lipid nanoparticle.

The type IV collagen alpha chains *COL4A3, COL4A4,* and *COL4A5* are expressed nearly exclusively in podocytes in the glomerulus. The three alpha chains trimerize into a mature α3-α4-α5 protomer and make up a critical structural component of the glomerular basement membrane (GBM) (**Figure 1B**). The GBM sits between glomerular endothelial cells and podocytes, all of which are important components of the glomerular filtration barrier (GFB).^8^ Mutations of *COL4A5* in in X-linked Alport syndrome (XLAS) or mutations in the other *COL4A* genes can disrupt the ability of these subunits to form protomers and led to a defective GBM and GFB (**Figure 1C**). When these fail, protein and blood cells leak into the urine leading to renal failure.

Since podocytes naturally express the three *COL4A* subunits, these cells are the presumptive target for AS molecular therapies. While podocytes reside in the glomerulus of the kidney where blood is filtered, they are not in direct contact with the blood. Instead, podocytes reside on the filtrate side of the GBM. This has significant implications for the ability to deliver *COL4A* nucleic acid therapies to these cells to treat AS.

The GFB is normally a significant barrier against any intravenous molecular therapy vector to penetrate into the kidney.^9,10^ The GFB is selectively permeable, with particle size and permeability being negatively correlated. Small proteins and particles such as inulin (∼5 kDa or 1.5 nm) are freely filtered through the GFB. In contrast, larger proteins and particles including albumin (∼3.55 nm or 69 kDa) are excluded by the GFB and are not normally filtered by the glomerulus.^11^ This poses a significant challenge to the development of gene and mRNA therapies using common molecular therapy vectors as these typically range in size from 25-200 nm in diameter (i.e. 25 nm adeno-associated virus (AAV), 100 nm adenovirus, 70-100 nm lipid nanoparticles (LNPs), 200 nm retroviruses).^9^

AAVs are currently the most popular vector for in vivo gene therapy by the intravenous (IV) route. They are popular because their small size allows them to penetrate many tissues from the blood after an intravenous injection. While they penetrate well into many tissues, they do not penetrate well into the kidney due to the GFB. Previous studies have shown that IV injection of different AAV serotypes in mice mediates zero to low transduction of cells in the glomerulus and even less in the parenchyma of the kidney^10,12^. Even when delivered locally, podocyte transduction by AAV vectors is generally weak to entirely absent.^10^

The small genome size of AAV also limits its use for Alport Syndrome. AAV vectors have a genome size of 4.7 kilobases.^13^ When viral genes are removed, this normally allows only 4.5 kilobases of exogenous DNA to be inserted. This is a problem for AS, since the sizes of the coding DNAs (cDNAs) for *COL4A3, COL4A4,* and *COL4A5* are all approximately 5,000 bases in length and exceed the normal capacity of AAV vectors.

Given these limitations, in this work, we instead tested the ability of LNPs to treat AS. We hypothesized that this might be feasible, since LNPs do not have specific nucleic acid size limitations, so the 5,000 base pair *COL4A* mRNAs could fit in these non-viral vectors. Furthermore, LNPs can co-package more than one mRNA. Therefore, it might be feasible to not just deliver one *COL4A* mRNA, but all three. While LNPs and mRNAs might be used to treat AS, one caveat is that nucleoside modified mRNAs delivered by LNPs mediate protein expression for only a brief period of time. mRNA expression typically peaks within 24 hours of injection and declines to baseline within a few days.^14^ We therefore hypothesized that any mRNA therapy for AS would require repeated dosing to maintain the therapy. While this was likely, this problem might be mitigated in part by the observation that type IV collagen in extracellular matrices can have half-lives over 100 days.^15^ We therefore hypothesized that this longer protein half-life might allow for longer intervals between mRNA treatments than with other more labile proteins. We also hypothesized that co-delivery of all three *COL4A* mRNAs might be more effective than delivering only one subunit, since co-delivery could maintain the 1:1:1 stoichiometry required for collagen IV α3-α4-α5 trimerization.

To test these hypotheses, experiments were performed to track mRNA transfection using LNPs *in vivo* in X-linked Alport syndrome (XLAS) mice.

## Results

### LNPs do not transfect the glomerulus when injected directly into the capsule of the kidney

We hypothesized that directly injecting LNPs into the parenchyma of the kidney might increase overall transfection in the kidney. To test this, LNP transfection tracking experiments were performed by packaging Cre recombinase mRNA and testing these in Cre reporter mice as in Hillestad et al.^16^ Transgenic mice used in this experiment harbor a membrane targeted tdTomato and GFP reporter cassette. The membrane targeted tdTomato fluorescent protein gene is flanked by loxP sites (floxed) upstream to a membrane targeted GFP gene. All cells in these mice constitutively express tdTomato unless transfected with LNPs carrying Cre recombinase mRNA. Once Cre recombinase is expressed in transfected cells, tdTomato is excised out of the reporter cassette, which activates constitutive expression of the downstream GFP.

To test the LNPs carrying Cre mRNA, one kidney in each reporter mouse was surgically exposed and the renal vasculature was secured with vascular microclamps to minimize fluid flow through the kidney. Each mouse was then injected with LNP-Cre mRNA directly into the kidney cortex through the capsule. Microclamps were removed after 5 minutes and then the incisions were sutured.

Under these conditions, kidneys injected with LNP-Cre mRNA exhibited robust GFP expression in long stretches of tubules and collecting ducts (**Supplemental Figure 1A**). Tubules were stained with Lotus Tetragonolobus Lectin (LTL). This showed transfection of these cells by the LNPs (**Supplemental Figure 1B**). In contrast, LNP-mRNA transfection was not observed in the glomerulus or in podocytes. This indicated that subcapsular injection was not likely to be a productive injection method to treat AS.

### LNP vector transfection in the kidney after IV injection is increased in Alport mice

We hypothesized that loss of GBM function in AS might allow LNPs to “leak” past the glomerular filtration barrier (**Figure 1C**). If so, this Alport pathology might be leveraged to reach podocytes by IV injection. To test this, the transgenic mice harboring the Cre-activated tdTomato/mGFP reporter cassette were crossed with X-linked Alport syndrome mice that have a nonsense mutation in exon 1 of the *Col4a5* gene.^17^ Healthier reporter mice and these XLAS Cre reporter mice were injected IV with LNPs carrying Cre mRNA. Four days after injection, organs were harvested and GFP expression was measured in the tissues using an IVIS imaging system (**Figure 2A**). Since cholesterol in LNPs targets LDL receptors, the liver was expected to absorb most of the IV dose of the LNPs.^18–20^ This was in fact the case where the livers of all animals showed significant expression of GFP (**Figure 2A and C**). No significant difference in GFP expression in the liver was observed between mice with XLAS or healthy mice (**Figure 2C)**. In contrast, the kidneys of XLAS reporter mice had 2-fold more GFP expression in the kidneys than in healthy reporter mice (**Figure 2A and B).** These data support our hypothesis that the pathology of Alport syndrome may increase vector transfection in the kidney.

**Figure 2:**
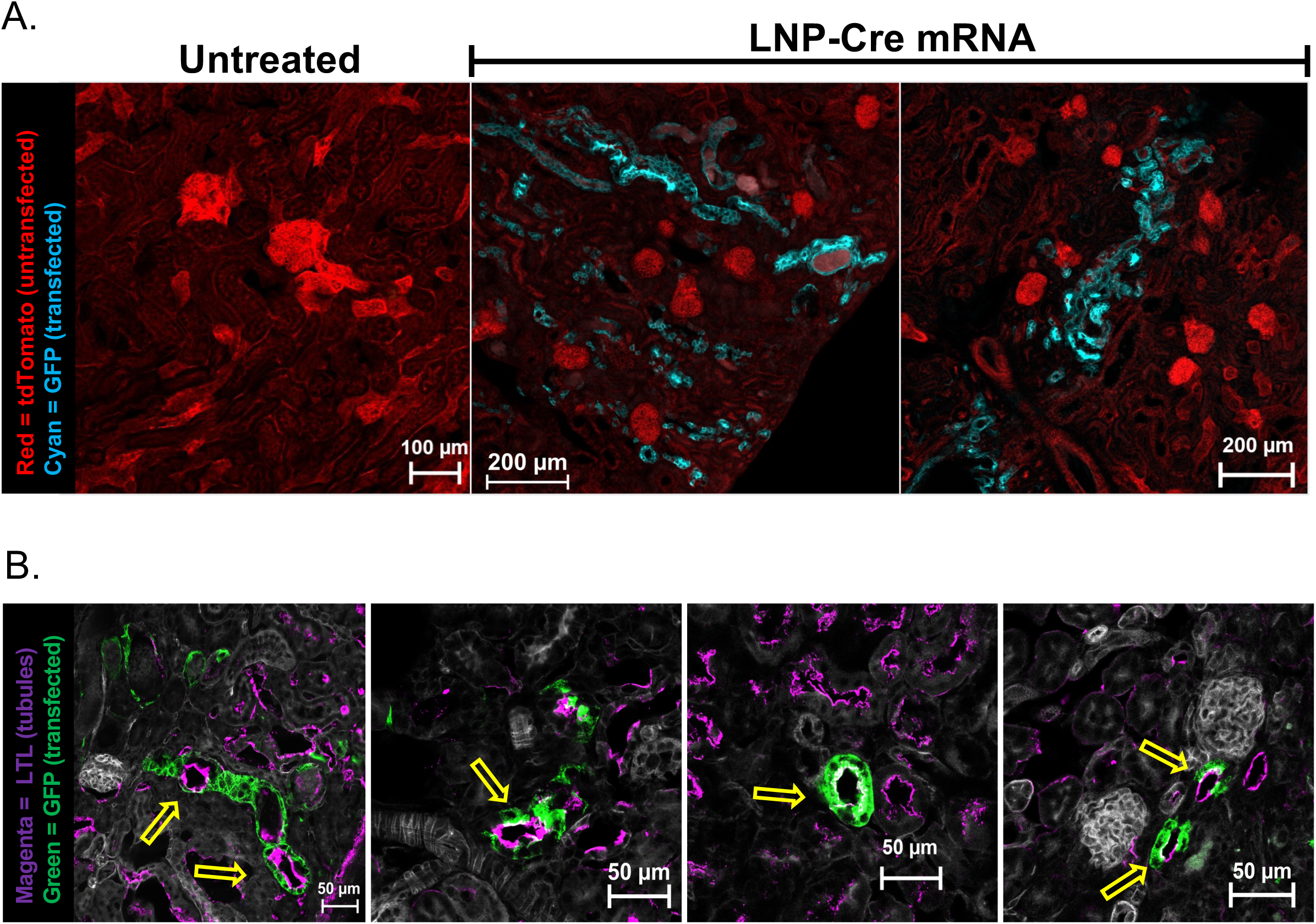
Comparison of LNP transfection in healthy and Alport mouse kidneys after single IV injection. **(A)** Healthy (left panels) and Alport (right panels) mouse kidneys and livers harvested 4 days after IV injection. GFP expression measured with IVIS imaging system. Top and bottom kidney panels represent two separate experiments; liver panel is representative of one experiment (n=6 mice total per group). LK, left kidney; RK, right kidney. **(B)** GFP expression in the kidneys, **(C)** and livers after a single IV injection, quantified as average radiance (photons/sec/cm^2^/sr)**. (D)** LNP transfection in the kidney increases with age and XLAS disease progression. Comparisons by unpaired t-test. ****p=<0.0001; NS, not significant. Data represents mean ± SD.

### The ability to transfect the kidney increases with disease progression

XLAS mice have normal kidney function at birth. After approximately 10 weeks of age, proteinuria increases as the GBM breaks down.^17^ To determine if AS disease progression affects the permeability of the GFB and transfection, progressively aged XLAS reporter mice were injected IV with LNPs with Cre mRNA and GFP expression was measured in their kidneys ex vivo using an IVIS imager (**Figure 2D**). Under these conditions, increasing LNP transfection was observed with increasing age in the XLAS animals.

### Intravenously delivered lipid nanoparticles transfect the glomerulus in Alport mice

Frozen tissue sections from LNP-Cre injected mouse kidneys were imaged using confocal microscopy to determine if the increased transfection seen in Alport kidneys was localized in the glomerulus. Little GFP expression was observed in healthy kidneys after IV Cre mRNA injections (**Figure 3A and B**). In contrast, many GFP-positive cells were observed in the glomeruli in Alport mice (**Figure 3C and D**). Most transfection was observed in the glomeruli of the XLAS reporter mice, but sparse transfection of other cell types was also observed. Tissue sections were counterstained with podocalyxin, a marker of podocytes. Colocalization analysis of podocalyxin and GFP demonstrated that in a given 2-dimensional slice, approximately 2.5% of podocalyxin-positive cells colocalized with GFP (**Supplemental Figure S2**).

**Figure 3:**
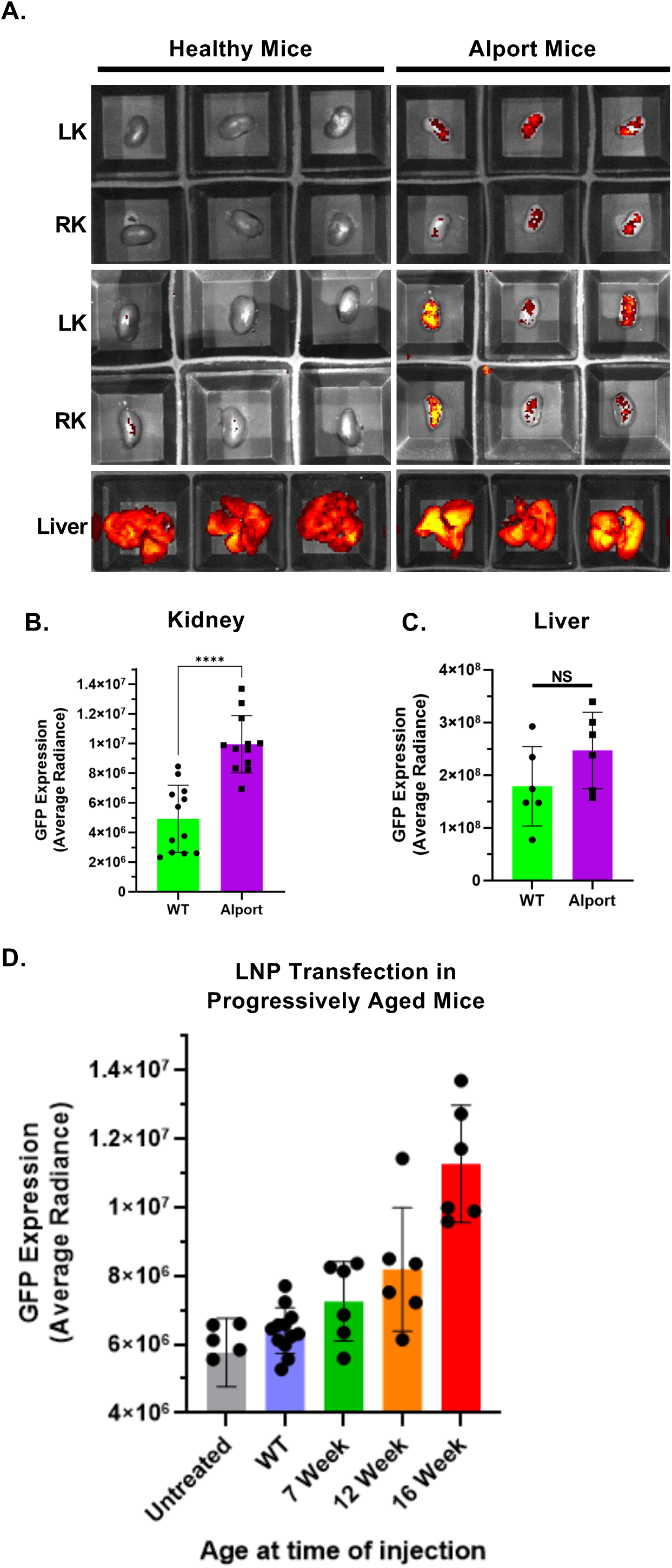
Comparison of LNP transfected cells in healthy and Alport kidneys after IV injection. **(A)** Composite tile scan of healthy (*Col4a5* WT) mouse kidney sections shows minimal LNP transfection in the kidney. Tile scans are composite images using a 20x objective. **(B)** Podocalyxin stained glomeruli shows rare LNP transfection. Taken with 40x water immersion objective. **(C)** Tile scan showing increased glomeruli transfection and GFP expression in Alport kidneys (yellow arrows). **(D)** Podocalyxin staining shows increased LNP transfection in the glomerulus in Alport kidneys. Taken with 40x water immersion objective.

### mRNA therapy administered after disease onset in aged XLAS mice does not significantly improve kidney function

These data demonstrated that LNP vectors could transfect the glomerulus and podocytes when delivered by the IV route and that the percent of cells transfected increased with increasing onset of proteinuria. While 2.5% podocyte transfection is relatively low, the fact that collagen IV protomers are secreted from cells suggested that these cells might be able to repair the GBM in a paracrine fashion.

Given these observations, we tested the ability of co-delivery and expression of all three collagen IV subunits by co-delivery of mRNAs expressing human *COL4A3, COL4A4,* and *COL4A5.* Aged 16-19 week old XLAS mice with already occurring proteinuria were treated IV with LNPs carrying the three *COL4A* mRNAs (LNP-COL4A3/4/5) (**Supplemental Figure 3**). These intravenous treatments were repeated every two weeks over 12 weeks for a total of 7 mRNA injections. Untreated XLAS mice of the same age were used as controls. The results presented here represent two separate experiments.

Urine and blood spots were collected from the aged mice and these were assayed for urine albumin, urine creatinine, and blood urea nitrogen (BUN). Spot urine collections vary in volume, so urinary albumin was normalized to urinary creatine to produce an albumin to creatinine ratio (ACR) for each sample (**Supplemental Figure 4**).

After LNP-mRNA treatment, ACRs trended lower in the aged XLAS mice when compared to untreated animals, however, the difference was not statistically significant (**Figure 4A**). Likewise, no significant differences in BUN levels were observed between treated and untreated mice (**Figure 4B**). Disease progression and survival was also similar in treated and untreated animals with ACRs significantly elevating in both groups by 12 weeks after the start of therapy. These data suggest that LNP-COL4A3/4/5 mRNA therapy is not effective once rampant disease progression is underway.

**Figure 4:**
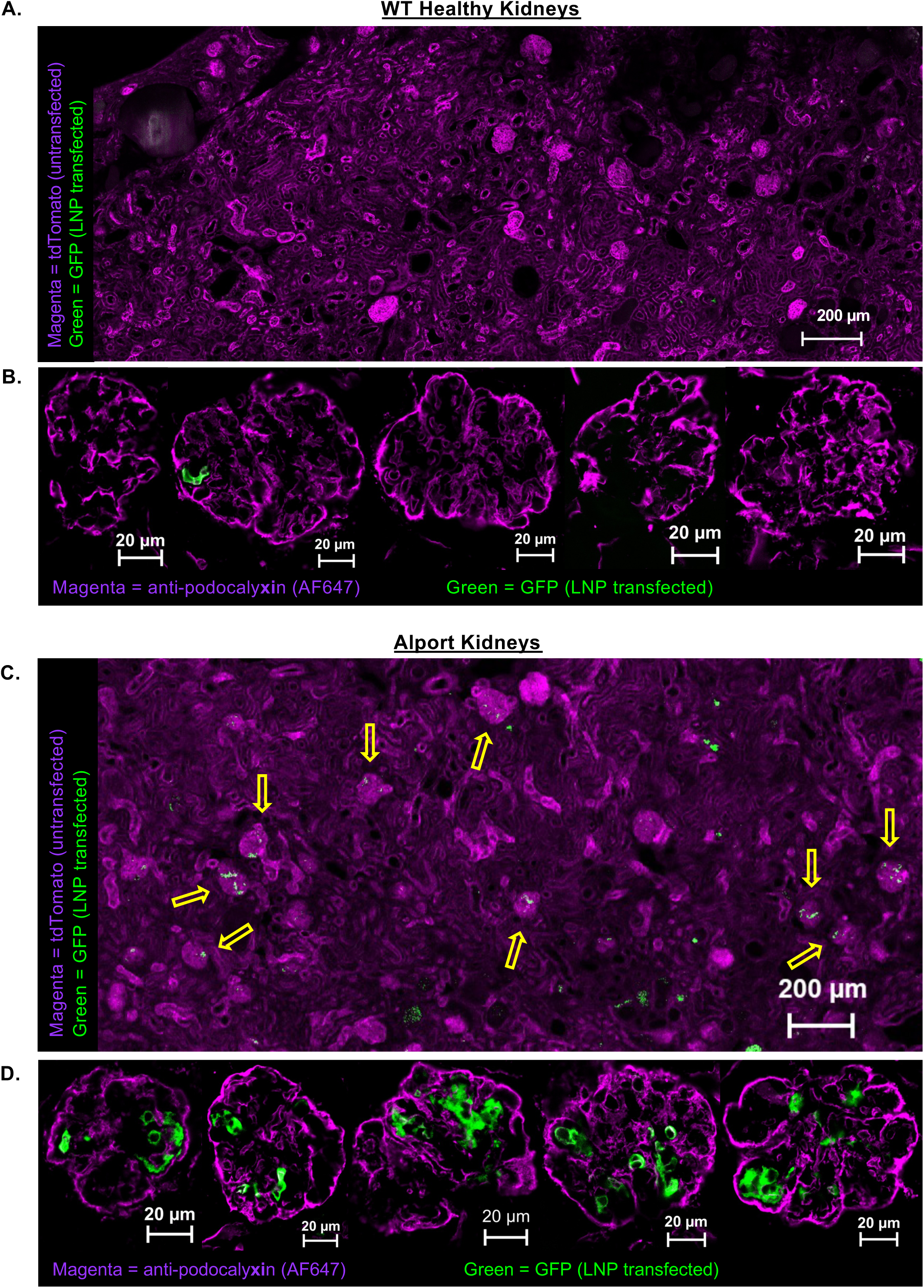
LNP-COL4A3/4/5 mRNA treatment after disease onset in aged XLAS mice. Mice aged 16-19 weeks at time of initial LNP-COL4A3/4/5 treatment. **(A)** Albumin to creatinine ratio (ACR) measurements. **(B)** Blood urea nitrogen (BUN) measurements. Black dotted lines represent values of healthy control mice. Comparisons by two-way ANOVA with Šídák’s multiple comparison test. *p= 0.0422. Data represents mean ± SEM.

### Early mRNA therapy administered before disease onset improves kidney function in Alport mice

Young 4-9 week old XLAS mice were next treated before manifestation of the disease. LNP-COL4A3/4/5 treatment was given IV by tail vein injection in two treatment series for a total of 7 doses of 13 µg mRNA per injection (**Figure 5A**). ACRs were comparable in all groups until week 6 after start of treatment. As the disease progressed and ACRs began to climb in untreated mice, LNP-COL4A3/4/5 mRNA demonstrated significantly lower ACRs and proteinuria and BUN than untreated XLAS mice at week 19 (**Figure 5B, C, D**). Importantly, this effect was not mediated by the LNPs themselves, as mice treated with empty LNPs containing no mRNA did not have the same reductions in urine albumin, ACRs, or BUN. In fact, ACRs and proteinuria were actually somewhat higher in mice receiving empty LNPs than in untreated control mice at weeks 15 and 17 and BUN was elevated at week 27.

**Figure 5:**
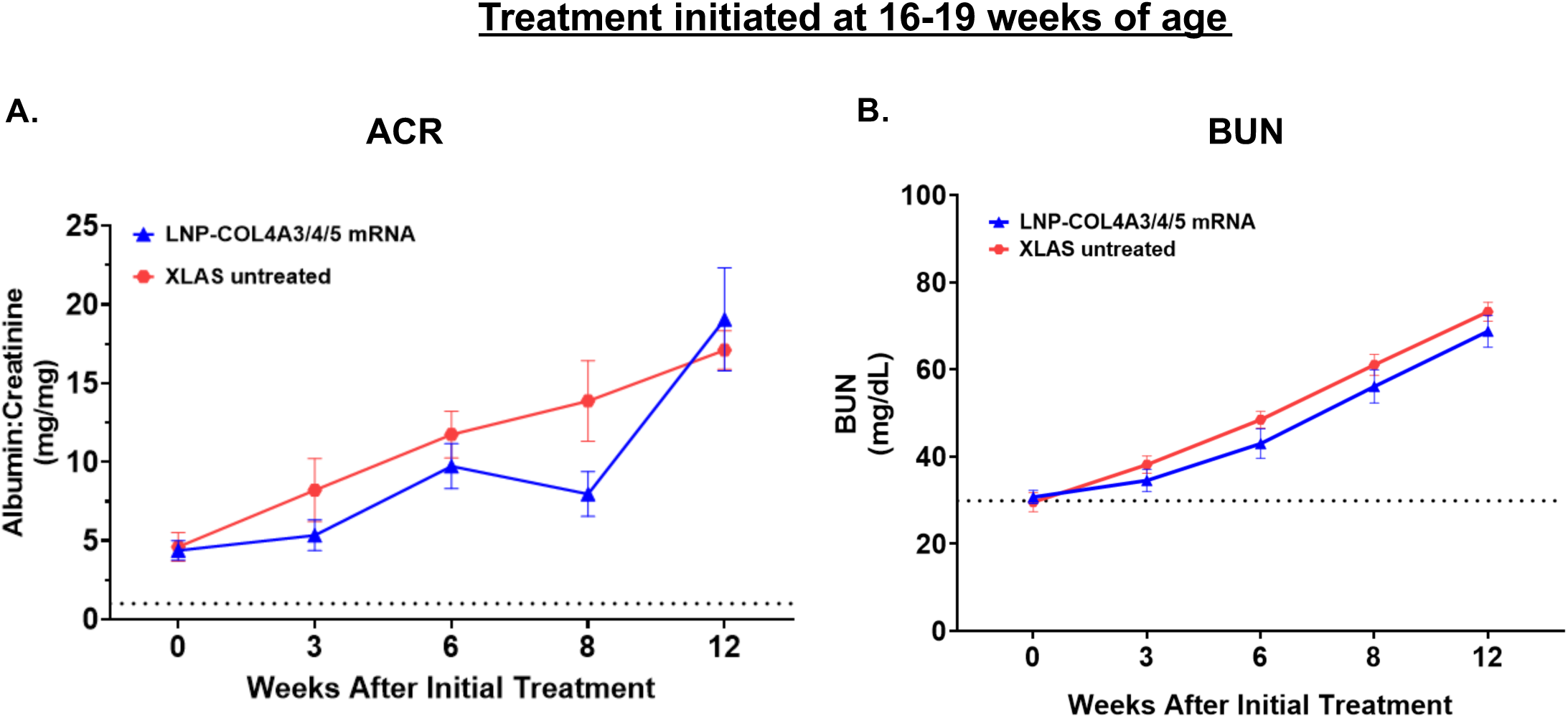
LNP-COL4A3/4/5 mRNA treatment prior to disease onset in XLAS mice. **(A)** Treatment schedule of LNP-COL4A3/4/5 and empty LNPs. **(B)** Urine albumin measurements, **(C)** Albumin to creatinine ratio (ACR) measurements, **(D)** Blood urea nitrogen (BUN) measurements at the indicated timepoints. Syringe icons represent a treatment date. Dotted black lines represent healthy control levels. Dotted red lines represent limit of detection. At week 27 and 30 the proportion of animals remaining in each group (color coded) are above the data points. Comparisons by two-way ANOVA with Šídák’s multiple comparisons test. ***p=0.0042; **p=≤0.0068; *p=≤0.0287. p-values represent the difference between LNP-COL4A3/4/5 mRNA treated and untreated XLAS mice. Data represents mean ± SEM.

### Efficacy is maintained during repeated mRNA therapy, but is lost when therapy ceases

Animals received their final treatment on week 19. Given that mRNA only expresses protein for a few days, it was expected that this mRNA-LNP therapy would be require continuous therapy to maintain protection. As expected, when the mRNA therapy was terminated, these animals lost control of their disease and urine albumin and ACRs increased, eventually matching those in untreated control animals (**Figure 5**).

## Discussion

In this study we tested the ability of LNP-mRNA therapy encoding human *COL4A3, COL4A4,* and *COL4A5* in equimolar ratios to mitigate Alport syndrome. We show that early mRNA intervention initiated before proteinuria occurs can delay disease progression, but intervention initiated after disease onset has limited effects. We show that the transient expression by mRNA maintains therapy if it is delivered repeatedly, but that this effect is lost when therapy ends.

The finding that LNP-COL4A3/4/5 therapy had the greatest benefit when initiated prior to disease onset aligns with previous evidence highlighting that early diagnosis and intervention with ACE/ARB drugs are critical to prolonging renal function in Alport patients.^21,22^ Podocytes are post-mitotic cells that have limited ability to regenerate.^23,24^ Therefore, it is likely that most strategies to treat AS, including by gene or mRNA therapy, will need to be administered early to significantly enhance renal survival.

Another outcome of this work is the observation that the increased permeability of the GFB associated with AS may fortuitously facilitate molecular therapy for this disease. Failure to provide all three collagen IV α-chain subunits in AS leads to a dysfunctional GBM and GFB. This is known to allow larger proteins and whole blood cells to pass the GFB and enter the urine, known as proteinuria and hematuria, respectively. It was unknown if this would affect filtration of very large, megaDalton molecular therapies. We found that overall LNP transfection in the kidney after IV injection was 2-fold higher in mice with X-linked Alport syndrome than healthy control mice. This result is important to consider, not only for vector delivery for Alport syndrome, but also for molecular therapy of other diseases that could also leverage increased permeability of the kidney. In fact, our group has previously induced proteinuria by administering lipopolysaccharides (LPS), a highly inflammatory bacterial toxin that induces proteinuria, prior to injecting AAV vectors IV as a strategy to enhance the transduction of tubule cells in the kidney.^25^

The treatment series used in our study was based off the dichotomy between achieving early intervention and the progressive leakiness of the GFB. In our study, the amount of LNP transfection in the kidney was positively correlated with age, disease progression, and GFB “leakiness”. We therefore decided to utilize an “early” and “late” treatment series to consider both points.

The mouse model of X-linked Alport syndrome used in this study contains a nonsense mutation in the *Col4a5* gene ablating the expression of one of the three alpha chains necessary for trimerization of a mature protomer^17^. Although other studies have postulated that delivery of only the α5 chain may be necessary to treat Alport syndrome, we hypothesized that delivery of all three alpha chains (α3-α4-α5) in equal molecular ratios would maintain the 1:1:1 ratio necessary for trimerization^26,27^. Despite only one of the collagen IV alpha chain genes being mutated in many (but not all) cases of Alport syndrome, lack of a single alpha chain (α5 in XLAS for example) results in the degradation of the other two alpha chains (α3, α4) despite their continued expression.^28,29^ Collagen IV networks are relatively stable and may not require constitutively high levels of expression to maintain the extracellular matrix. Therefore, exogenous delivery of *COL4A5* nucleic acids alone may exceed the stoichiometry of the other two alpha chains that are still endogenously expressed.

Additionally, we hypothesized that providing the α3, α4, and α5 chains would allow expression in other cell types such as glomerular endothelial cells or hepatocytes to deliver type IV collagen *in trans*. It has been thought that glomerular endothelial cells could potentially help repair the GBM, but previous studies have shown reintroducing a single missing alpha chain in those cells does not rescue the disease phenotype in a drug inducible expression model as was achieved by reintroduction of the same gene in podocytes that still express the other two collagen IV alpha chains^27,30^. Suggesting that if glomerular endothelial cells, or other cells, could facilitate a reparative effect they would need to express all three alpha chains within the same cell. Another benefit of our strategy to deliver all three alpha chains in the same LNP vector (α3-α4-α5), is this therapeutic could be adapted to autosomal dominant and autosomal recessive disease where the α3 or α4 chain is mutated.

An unexpected result of these experiments was the increase in ACR and BUN levels in empty LNP treated mice (**Figure 5**). This would indicate that the lipid components themselves may accelerate the progression of Alport syndrome. While the exact mechanism of this effect is unknown, excess lipid accumulation in podocytes is a known driver of lipotoxicity in CKD and Alport syndrome^31–35^. This could dampen the net therapeutic effect of LNP-COL4A3/4/5. It is currently unclear if this is being caused by single or multiple lipid components.

A major limitation with mRNA therapeutics is that the duration of expression is short lived compared to DNA based expression systems. We hypothesized that LNP-mRNA therapeutics could be utilized in Alport syndrome because type IV collagen is a stable protein with a relatively low turnover rate, and therefore short bursts of mRNA expression could provide longer lasting therapy than is typically achieved with mRNA.^15^ In our study, the therapeutic effect of LNP-COL4A3/4/5 when given before disease onset was durable until the treatment ceased (**Figure 5**). Eight weeks after the final injection, the urine albumin and ACRs of the LNP-mRNA treated mice deteriorated but were still below the levels of untreated control mice with XLAS at week 27 (**Figure 5B,C**). This indicated that the LNP-mRNA therapeutic effect outlasted the short duration of mRNA expression. Although mRNA therapeutics will need repeat dosing, this study shows that the time between doses can be extended to reduce the frequency needed to see an effect, which would be a substantial benefit if translated to the clinic. Our study demonstrates that LNP-mRNA therapeutics could be utilized to treat genetic diseases where the missing protein in question is a stable, low-turnover protein.

Although IV injections are simple to perform, this was a major limitation of our study as an effect of using mouse models. A consequence of systemic delivery by IV injection was a significant amount of off-target transfection in the liver (**Figure 2A,C**). A result that was expected given that LNPs are largely liver tropic due to the primary receptor utilization of low-density lipoprotein receptor (LDLR) which is abundant in the liver. Another inefficiency of IV injection in our study was a low percentage of transfected podocytes. It is important to note that our LNP-Cre tracking experiments were performed by a single IV injection, and the animals treated in the therapeutic efficacy study were treated a total of 7 times, likely leading to an increase in the number of total glomerular cells transfected. Previous studies using IV AAV at relatively high doses to treat glomerular disease also reported low podocyte transduction (∼10%) with a smaller diameter viral vector than our LNPs. Despite low transduction, they were able to show meaningful improvements in their disease model after treatment, indicating that therapeutic efficacy in some models can be achieved even if large proportions of the podocytes are not directly targeted.^12^

Advanced methods could be adapted for delivering LNPs or other vectors directly to the renal blood supply, such as arterial catheterization or stenting, renal perfusion systems, or ultrasound guided injection methods.^36–38^ Utilizing more precise methods, which are more readily applicable to larger animal models or humans in a clinical setting, could significantly decrease off-target transfection and greatly increase the delivery of these vectors to the kidney, and more specifically the glomerulus.

In conclusion, LNPs were able to transfect the kidneys and glomeruli of Alport mice more effectively than healthy control mice. Indicating that the same mechanisms that cause proteinuria and hematuria in Alport patients can aid in delivery of molecular therapies compared to non-permissive healthy kidneys. Additionally, intravenous injections of LNP-mRNA encoding *COL4A3, COL4A4,* and *COL4A5* were able to improve kidney function in a mouse model of XLAS.

## Materials and Methods

### Plasmid construction and *in vitro* transcription (IVT)

Coding DNA for each gene was cloned into an IVT plasmid containing a T7 promoter and a poly-adenosine tract. Human sequences were used for all collagen genes. The plasmids were then linearized by restriction digest with ApoI and purified using Phenol/Chloroform/Isoamyl alcohol extraction followed by two rounds of chloroform back extraction to remove residual phenol, followed by ethanol precipitation. These linearized plasmids served as the template for IVT using a T7 RNA synthesis kit (New England Biolabs HiScribe E2040S). All mRNAs were synthesized using N1-Methylpseudouridine-5’-Triphosphate (m1ΨTP, Trilink N-1081). A synthetic capping reagent (Trilink CleanCap reagent AG N-7113) was used for co-transcriptional capping of the mRNAs. DNase I treatment was performed to remove the DNA template from the synthesized mRNA, then mRNAs were purified by column purification (NEB Monarch RNA Cleanup Kit T2050). mRNA was then denatured at 70°C for 10 minutes in RNA denaturing loading dye (NEB 2x RNA loading dye) and quality was assessed by 1% agarose gel (**Supplemental figures 3 & 5)** and nanodrop.

### Lipid Nanoparticle Assembly

Lipid nanoparticles were formulated using Cholesterol (Sigma C3045), 1,2-dimyristoyl-rac-glycero-3-methoxypolyethylene glycol-2000 (DMG-PEG2000, Avanti polar lipids 880151P), 1,2-distearoyl-sn-glycero-3-phosphocholine (DSPC, Avanti polar lipids 850365P), and dilinoleylmethyl-4-dimethylaminobutyrate (D-Lin-MC3-DMA or “MC3”, MedChemExpress HY-112251). Lipids were resuspended in 100% Ethanol at a molar ratio of 10:48:40:2 (DSPC:Cholesterol:MC3:DMG-PEG2000). LNP assembly was performed using a NanoAssemblr Ignite (Cytiva) microfluidics system. The mRNA was suspended in 100 mM citrate buffer at pH 4 to facilitate ionization of the MC3 lipid when mixed with the lipids in ethanol. Lipids in ethanol and mRNA in citrate buffer were mixed at a ratio of 3:1, respectively, and at a total flow rate of 12 mL/min. The assembled LNPs were then diluted in PBS (without calcium or magnesium) 40x, followed by reconcentration with Amicon 10,000 MWCO centrifugal filters (Millipore Sigma UFC901096). The concentrated LNPs were then filtered with a 0.2 µM syringe filter (Pall Acrodisc 4602). Size (avg: ∼80nm diameter), polydispersity index (<0.1), particle concentration (approximately 1.5×10^13^ particles/mL average), and zeta potential (∼ -3 to - 5 mV) were measured using a Zetasizer Advance Ultra Red (Malvern Panalytical ZSU3305). mRNA encapsulation efficiency and total encapsulated mRNA was quantified using a Quant-iT RiboGreen RNA assay kit (Invitrogen R11490).

### *In Vivo* Experiments

All animal experiments were approved by the Mayo Clinic Institutional Animal Care and Use Committee. Animals were housed at Mayo Clinic facilities and all experiments followed guidance of the Public Health Service Animal Welfare Policy, Animal Welfare Act, and the NIH guidelines in the *Guide for the Care and Use of Laboratory Animals*.

### LNP vector tracking experiments

STOCK Gt(ROSA)26Sortm4(ACTB-tdTomato,-EGFP)Luo/J (JAX strain# 007576) mice (mT/mG mice) were bred with B6.Cg-*Col4a5*^tm1Yseg^/J (JAX strain# 006183) mice to create a XLAS hybrid reporter mouse model (XLAS reporter mice). The resulting offspring are heterozygous for the membrane bound tdTomato/mGFP reporter cassette and are genotyped to determine *Col4a5* mutants (XLAS) from *Col4a5* WT mice (healthy). These mice were injected with 100 µL of LNP-Cre mRNA (14 µg per mouse) by intravenous tail vein injection. Three mice for each group were injected in each experiment and the experiment was repeated once for a total n=6 per group. Four days after injection, organs were removed and imaged using a Lumina S5 IVIS imager (Revvity) to measure GFP expression. After imaging, organs were fixed and frozen using the “tissue preservation, freezing, and sectioning” method described below. Once microscope slides were prepared, a hydrophobic barrier pen was used around each sample, then tissues were blocked with blocking buffer (1% BSA in PBS) for 2 hours at room temperature. The blocking buffer was then removed and replaced with a polyclonal goat anti-mouse podocalyxin primary antibody diluted in blocking buffer at a 1:200 dilution (R&D Systems cat# AF1556) and incubated overnight in a humidified chamber, protected from light, at 4°C. The primary antibody was then aspirated, and slides were washed 5 times in PBS before adding the donkey anti-goat IgG-AF647 secondary antibody in blocking buffer at a 1:10,000 dilution, and incubated at room temperature for 1 hour. Slides were washed again 5 times in PBS and excess liquid removed before application of VectaShield antifade reagent (Vector Labs cat# H-1000). Stained tissue sections were then imaged by confocal microscopy (Zeiss LSM 780). Colocalization analysis was performed using ImageJ JACoP.

For subcapsular injection experiments, STOCK Gt(ROSA)26Sortm4(ACTB-tdTomato,-EGFP)Luo/J (JAX strain# 007576) mice (mT/mG) were used. Each mouse was anesthetized by isoflurane inhalation. The site of incision was shaved and prepared with 3 rounds of alternating iodine and ethanol wipes. The right kidney was surgically isolated, and renal vasculature was clamped using microvessel clamps. Using a 31-gauge insulin syringe (BD) 50 µL of LNP-Cre mRNA (9.87 µg mRNA n=2) was injected into the cortex of the kidney. After injection, the microvessel clamps and injection needle remained in place for 5 minutes before releasing the clams and closing the incision. Kidneys were harvested 7 days after injection. Tissues were processed and prepared according to the “tissue preservation, freezing, and sectioning” method described below.

### LNP transfection in progressively aged mice experiment

XLAS-reporter mice were injected with LNP-Cre mRNA by IV tail vein injection (14 µg mRNA per mouse) at 7, 12, and 16 weeks of age (n=3 per group). Untreated mice (n=3) and age matched healthy mice were used as control mice (n=2 per age group for total n=6). Four days after injection, animals were euthanized by CO_2_ inhalation, then kidneys were harvested and GFP expression was measured using an S5 Lumina IVIS imager.

### Tissue preservation, freezing, and sectioning

Animals were euthanized by CO_2_ inhalation and organs were moved to ice cold 4% paraformaldehyde (PFA) and transferred to 4°C overnight, protected from light. Then PFA was removed and replaced with 15% w/v sucrose in PBS for 6 hours (or until organs sink) then removed and placed in 30% w/v sucrose in PBS overnight until tissues sank. All sucrose steps were incubated at 4°C. Organs were then rinsed in PBS and flash frozen in optimal cutting temperature compound (OCT) using dry ice cooled isopentanes and stored at -80°C. Frozen tissues were sectioned using a cryostat in 18 µm slices and mounted on glass slides treated with HistoGrip (Thermo Fisher cat# 008050).

### Therapeutic efficacy experiment in X-linked Alport mice

For studies treating young mice prior to disease onset (**Figure 5**), B6.Cg-*Col4a5*^tm1Yseg^/J mice (JAX strain# 006183) (XLAS mice) 4-9 weeks of age were used. Before the first treatment, urine and serum were collected for baseline urine albumin, creatinine, and BUN measurements. XLAS mice were injected IV by tail vein with either empty LNPs (n=10) or LNP-COL4A3/4/5 mRNA (n=11) at week 0 (study start), 2, 4, 13, 15, 17, and 19. Each animal received 13 µg mRNA per injection for LNP-COL4A3/4/5 treatment, or an equivalent volume of empty LNPs. Untreated XLAS mice were also included as a reference for disease progression in the absence of intervention (n=10). Wild type *Col4a5* mice were also included as a reference for healthy levels of each measurement type (n=10). Urine was collected periodically as described below at weeks 0, 3, 6, 12, 15, 17, 19, 27, and 30. Blood was collected less frequently at weeks 0, 12, 19, 27, and 30 to minimize the risk associated with blood draws on the animal’s health. Measurement of kidney function was performed as described later in the methods section. Two independent experiments were performed.

For studies using aged mice treated after disease onset (**Figure 4**), male XLAS mice 16-19 weeks of age were used. XLAS males were either treated with LNP-COL4A3/4/5 mRNA (n=11) or untreated controls (n=11). LNPs were injected (13 µg mRNA) into the tail vein of each mouse biweekly from week 0 (study start) to week 12 for a total of 7 injections. Urine and blood were collected at weeks 0, 3, 6, 8, and 12. Urine albumin, urine creatinine, and BUN levels were measured as described later in the methods section. Two independent experiments were performed.

### Kidney Function Measurements

Urine collection was performed by manual restraint and collection of the voided urine into a 1.5 mL microfuge tube. Urine albumin and creatinine content were measured using QuantiChrom albumin (BioAssay Systems DIAG-250) and QuantiChrom creatinine assay kits (BioAssay Systems DICT-500). Blood draws were performed by cheek bleed and collection into BD microtainer amber collection tubes which were used to isolate serum. Serum was used to measure BUN levels using QuantiChrom urea (BioAssay Systems DIUR-100).

### Statistical analysis

Statistical analysis for this study was performed using GraphPad Prism version 10.4.1. Information regarding specific statistical tests used is in each figure legend.

## Acknowledgements

B.J.P. was supported in part by the Mayo Clinic Virology and Gene Therapy T32 training grant 5T32AI132165. This work was supported by Mayo Clinic Ventures Innovation Program Funding Request. Some figures for this manuscript were created using https://BioRender.com.

## Data availability Statement

Reasonable requests for materials, reagents, and data can be made by contacting the corresponding author.

---

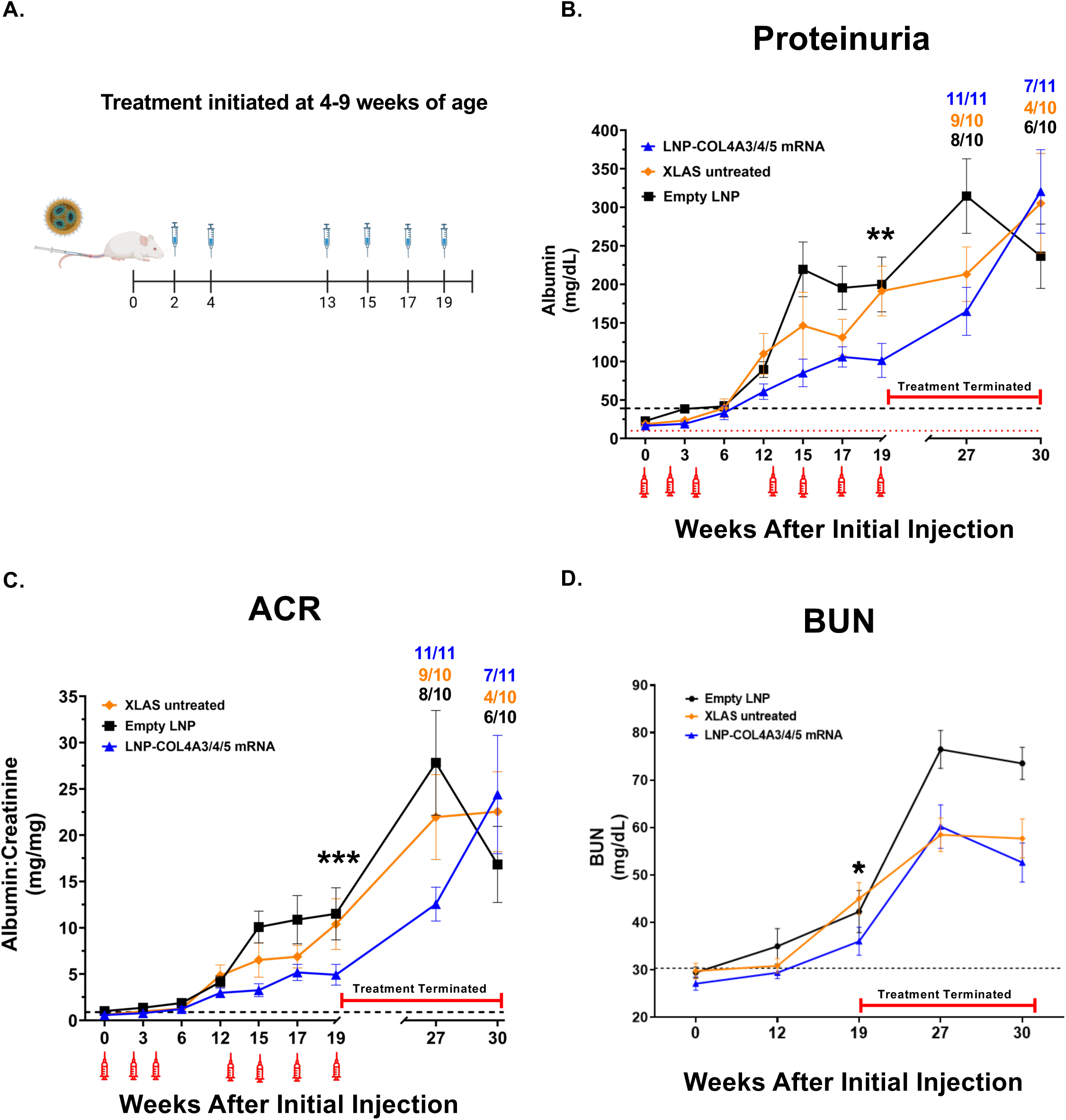

