## Supplemental Figures for "mRNA Therapy for Alport Syndrome"

**Figure S1**

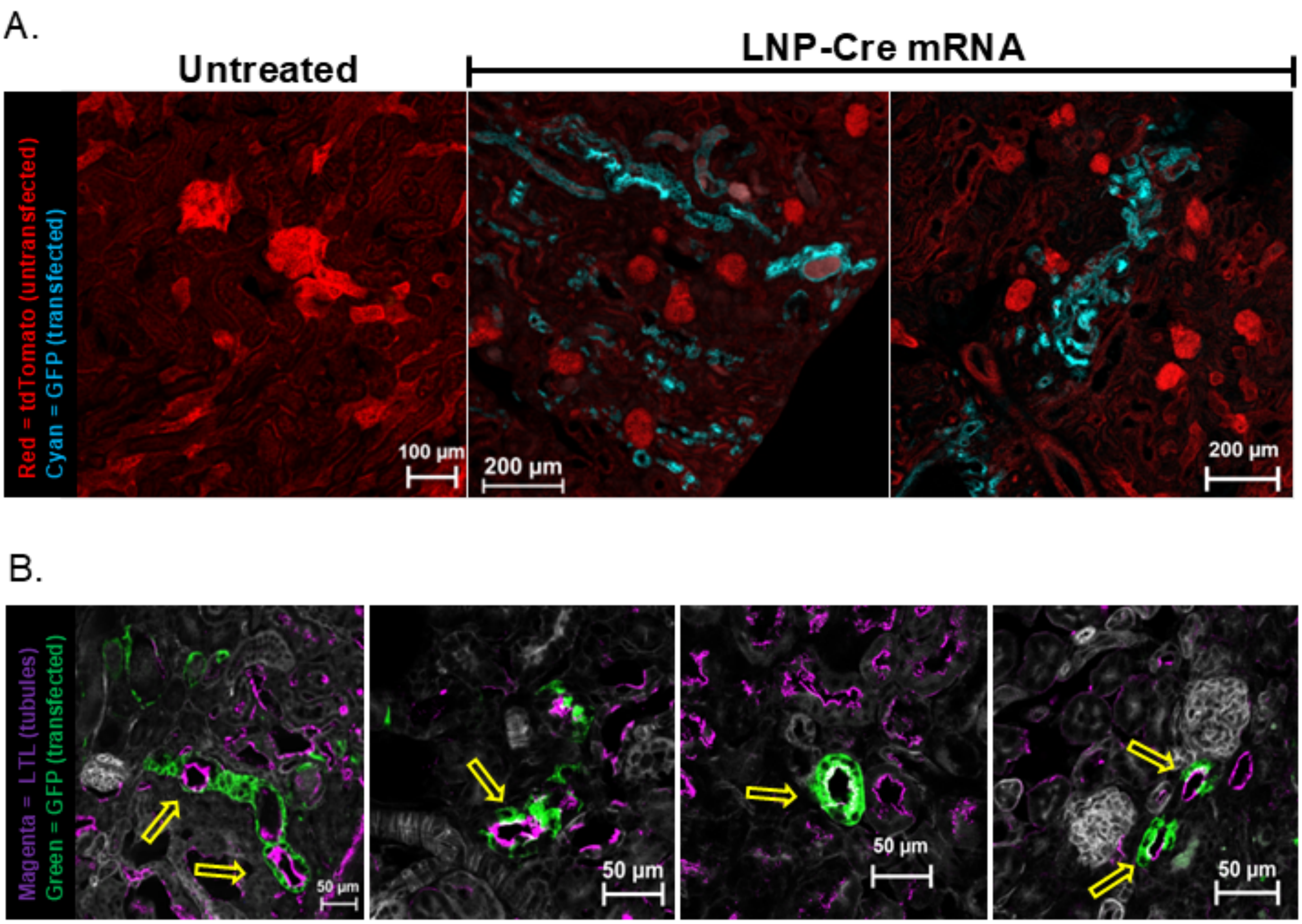


**Figure S1: LNP vector tracking after injection of LNPs by subcapsular injection. (A)** Representative kidney tissue section images of untreated and LNP-Cre mRNA injected kidneys. Untransfected tdTomato cells are red, LNP transfected GFP expressing cells are cyan. **(B)** LTL staining of LNP-Cre mRNA injected kidneys was performed to verify colocalization with GFP positive cells (yellow arrows). LTL staining in magenta, GFP positive transfected cells in green.

**Figure S2.**

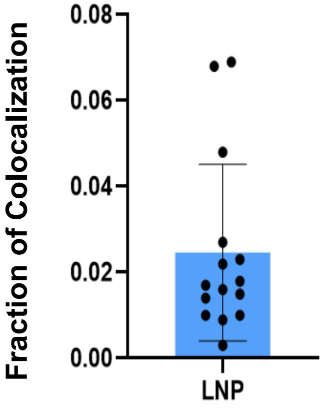


**Figure S2:** Colocalization analysis shows that an average of ~2.5% of podocytes were transfected per glomerulus (n=15). Analysis performed with ImageJ JACoP. Data represents mean ± SD.

**Figure S3.**

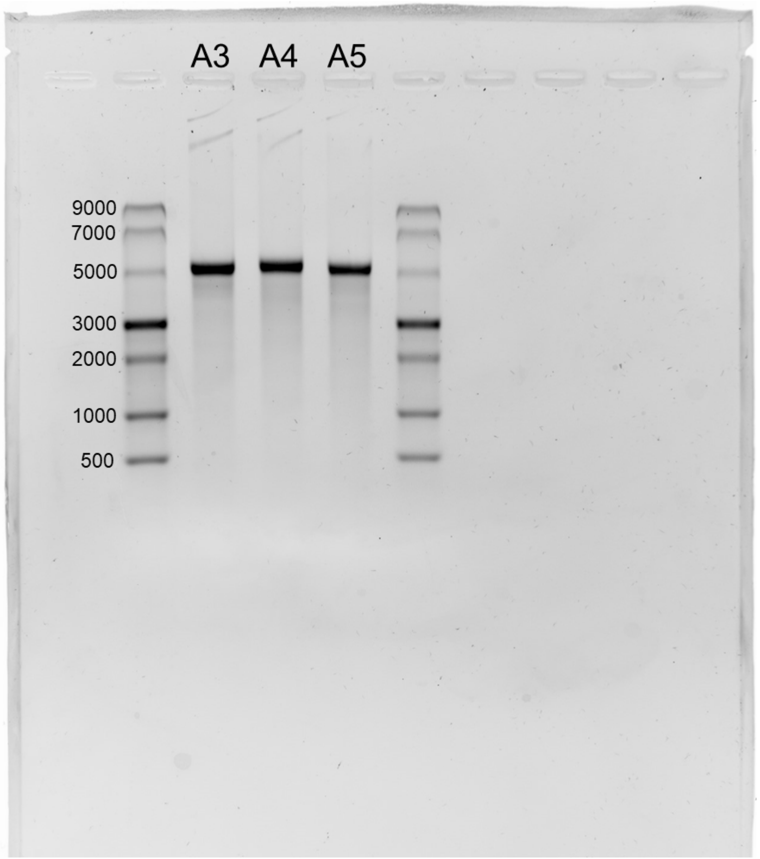


**Figure S3:** COL4A3, COL4A4, COL4A5 mRNA ran on agarose gel after IVT and purification.

**Figure S4**

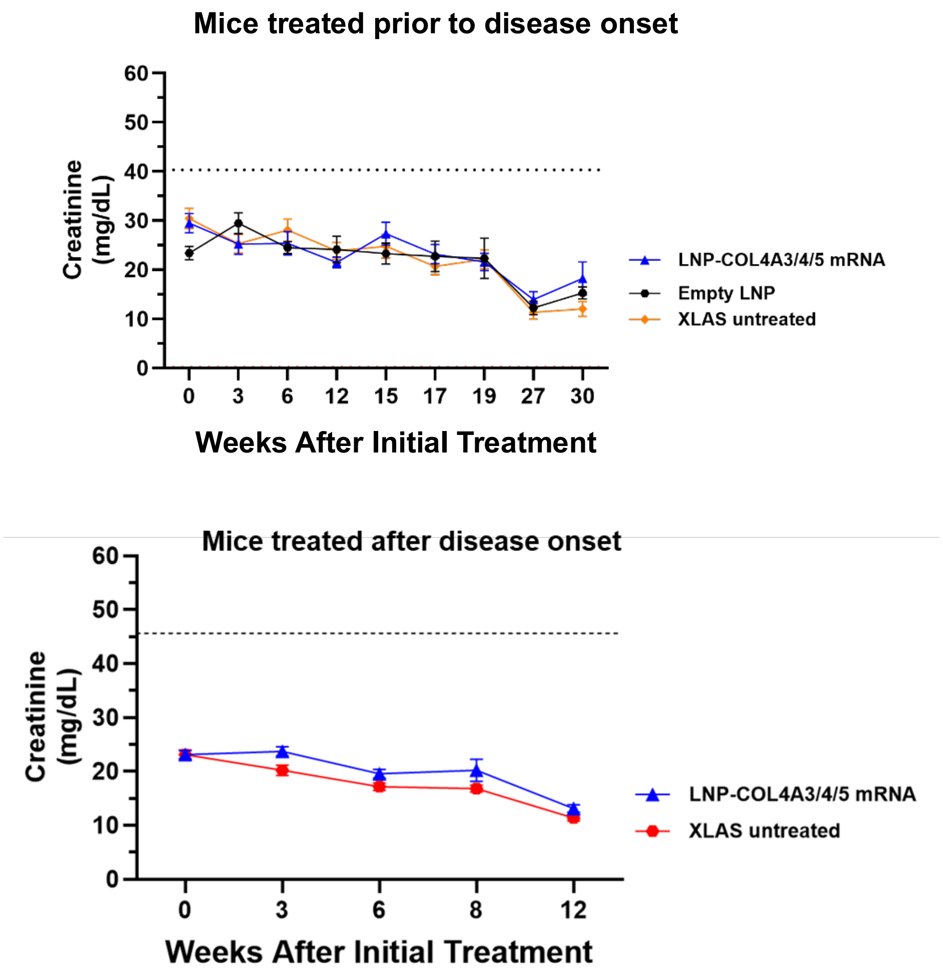


**Figure S4:** Urine creatinine measurements for each timepoint used for calculating ACRs. Black dotted line represents creatinine levels of healthy controls. Top graph represents data from the experiment where young mice were treated prior to disease onset. Bottom graph represents data from the experiment where older mice were treated after disease onset.

**Figure S5.**

Cre mRNA (1301 bp)

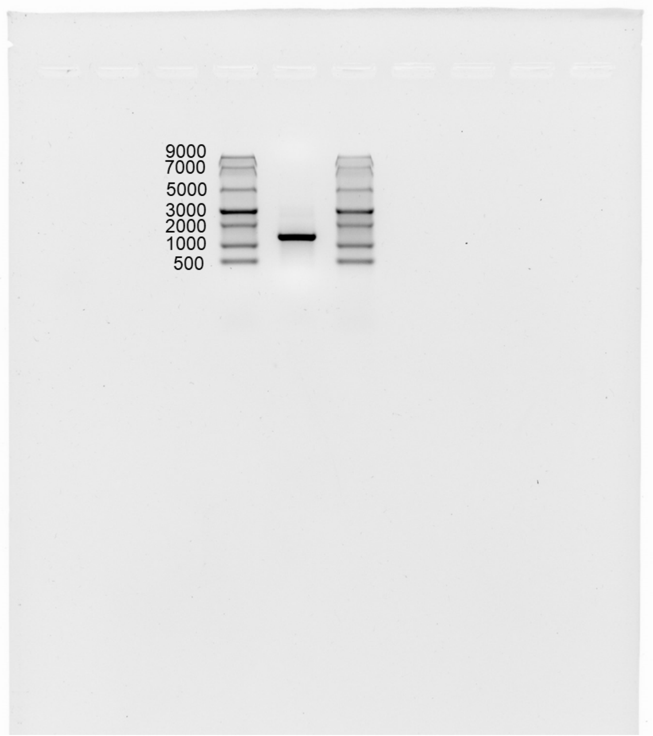


**Figure S5:** Agarose gel image of purified Cre mRNA after *in vitro* transcription (IVT). Cre mRNA is ran between both ssRNA ladders (NEB ssRNA ladder).
